# The effect of presentation level on spectrotemporal modulation detection

**DOI:** 10.1101/450957

**Authors:** Sara Magits, Arturo Moncada-Torres, Lieselot Van Deun, Jan Wouters, Astrid van Wieringen, Tom Francart

## Abstract

The understanding of speech in noise relies (at least partially) on spectrotemporal modulation sensitivity. This sensitivity can be measured by spectral ripple tests, which can be administered at different presentation levels. However, it is not known how presentation level affects spectrotemporal modulation thresholds. In this work, we present behavioral data for normal-hearing adults which show that at higher ripple densities (2 and 4 ripples/oct), increasing presentation level led to worse discrimination thresholds. Results of a computational model suggested that the higher thresholds could be explained by a worsening of the spectrotemporal representation in the auditory nerve due to broadening of cochlear filters and neural activity saturation. Our results demonstrate the importance of taking presentation level into account when administering spectrotemporal modulation detection tests.

## 1. Introduction

Complex acoustic signals such as speech are characterized by a combination of spectral and temporal modulations. Speech understanding relies (at least partially) on the ability to detect and discriminate these modulations. In other words, it relies on an individual’s spectrotemporal modulation sensitivity (Supin et al., 1997). This can be assessed by two categories of tests: *spectral ripple discrimination* tests and *spectral/spectrotemporal modulation detection* (SMD/STMD, respectively) tests. There are many varieties of these. However, in this paper we focus on an SMD/STMD test where participants are asked to discriminate between a modulated and unmodulated stimulus. The modulation detection threshold is usually defined as the minimal peak-to-valley ratio or modulation index at which the participant can discriminate between the two stimuli (e.g., Bernstein et al., 2013).

It has been shown that SMD/STMD thresholds are correlated with different measures of speech perception in quiet and in noise (Anderson et al., 2012; Mehraei et al., 2014; Davies-Venn et al., 2015; Croghan and Smith, 2018). Additionally, SMD/STMD thresholds can provide a non-linguistic measure of spectral/spectrotemporal sensitivity without the confounding factor of language knowledge that plays a role in standardized tests (e.g., speech audiometry, Gifford et al., 2014; Davies-Venn et al., 2015; Choi et al., 2016). This has motivated their use for a variety of purposes. For example, STMD paradigms have been used to explore perceptual learning mechanisms in the auditory system (Sabin et al., 2012). SMD/STMD tests have also used spectral/spectrotemporal resolution successfully as an outcome measure in different fields of audiological research: prediction of speech understanding in noise of hearing-aid users (Bernstein et al., 2016), assessment of cochlear implant candidacy, parameter fitting, and new sound processing strategies (Langner et al., 2017; Choi et al., 2016; Croghan and Smith, 2018; Zheng et al., 2017), evaluation of bimodal hearing benefit (Zhang et al., 2013), and music perception (Choi et al., 2018).

Although these tests are used mostly in audiological research, to our knowledge no studies have evaluated how presentation level affects SMD/STMD thresholds. This is relevant because SMD/STMD thresholds might be negatively affected by the broadening of the auditory filters with increasing presentation level (Glasberg and Moore, 2000). Taking the effect of level into account is crucial when administering SMD/STMD tests in a research environment, in (potential) clinical practice, and even more in test situations where it cannot be controlled strictly (e.g., home-based computerized rehabilitation programs). Furthermore, we need to understand this effect to be able to make a fair comparison of behavioral SMD/STMD results obtained at different presentation levels within and across studies.

The goal of this work was to explore how presentation level affects SMD/STMD thresholds for young adult NH participants. Specifically, we focused on the STMD test, since spectrotemporally modulated (i.e., moving spectral ripple) stimuli have been suggested to provide a better representation of speech (Won et al., 2015) than stimuli measuring sensitivity to only spectral (i.e., rippled, Litvak et al., 2007; Saoji et al., 2009) or temporal modulation. Additionally, STMD tests prevent participants from having access to phase cues by using low rate temporal modulation (Bernstein et al., 2013). Furthermore, we used a biologically inspired model of peripheral processing up to the auditory nerve (AN) to help us interpret the behavioral results, to study the contribution of peripheral information to spectrotemporal sensitivity, and to generate STMD threshold predictions.

## 2. Behavioral Measurements

### 2.1. Materials & Methods

#### 2.1.1. Participants

Ten participants (1 male, 9 female, median age 23.5 years, age range 21–29 years) took part. They had audiometric thresholds ≤ 20 dB HL at all octave frequencies from 125 to 8000 Hz. Written informed consent was obtained. The study was approved by the Ethics Committee of the University Hospitals Leuven (approval no. B322201731501).

#### 2.1.2. Equipment

Measurements were performed in a double-walled sound-attenuating booth. Stimuli were played from a computer via an RME Hammerfall DSP Multiface II sound card and presented to the participants through Sennheiser HDA 200 headphones using APEX 3 (Francart et al., 2008).

#### 2.1.3. Stimuli

We used the spectrotemporally modulated stimuli described by Kowalski et al. (1996) and Chi et al. (1999). These were 500-ms long (including 20-ms onset and offset cosine ramps) and were generated with a sampling frequency of 44100 Hz and 16-bit resolution using MATLAB (Mathworks, Natick, MA).

The spectral modulation was achieved as follows. The spectrum of the ripple stimulus (the “carrier”) consisted of 4000 random-phase tones equally spaced along the (logarithmic) frequency axis from 354 to 5656 Hz. The amplitudes of the individual components were adjusted to form a sinusoidally shaped spectrum around a flat base. The amplitude of the ripple was defined as the modulation depth *m*. The initial phase of the ripple Φ was defined relative to a sine wave starting at the low-frequency edge. Its value was set using 50 different selections of random phases between 0 and 2π to prevent participants from using phase differences as a cue. The ripple density was defined as Ω (with values of 0.5, 2, and 4 ripples/oct). The mathematical expression for the static ripple is given in Eq. 1

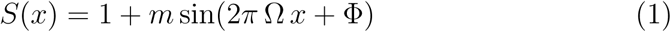

where *x* is the position on the logarithmic frequency axis (in octaves), which was defined as 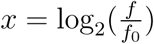 with *f* being the component tone frequency and *f*_0_ the low-frequency edge. Notice that when *m* = 0, the resulting profile is a flat spectrum.

The temporal modulation was achieved by moving the static ripple downwards along the frequency axis at a constant velocity ω (defined as the number of ripple per second passing the low-frequency edge of the spectrum). The value of ω was 4 Hz. The complete mathematical expression for the spectrotemporal modulated stimuli is given in Eq. 2, where *t* is time.

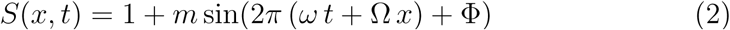

In order to make our results comparable to those of previous studies, we report the modulation depth *m* as 20 log_10_(*m*) (i.e., in dB). The *reference stimulus* was unmodulated (i.e., 20 log_10_ (*m*) = − ∞ dB), whereas the modulation depth of the *target stimulus* was varied adaptively (Sec. 2.1.4). Figure 1 shows spectrograms of the reference stimulus and two example target stimuli (20 log_10_(*m*) = −6 dB and 20 log_10_(*m*) = 0 dB).

**Figure 1:**
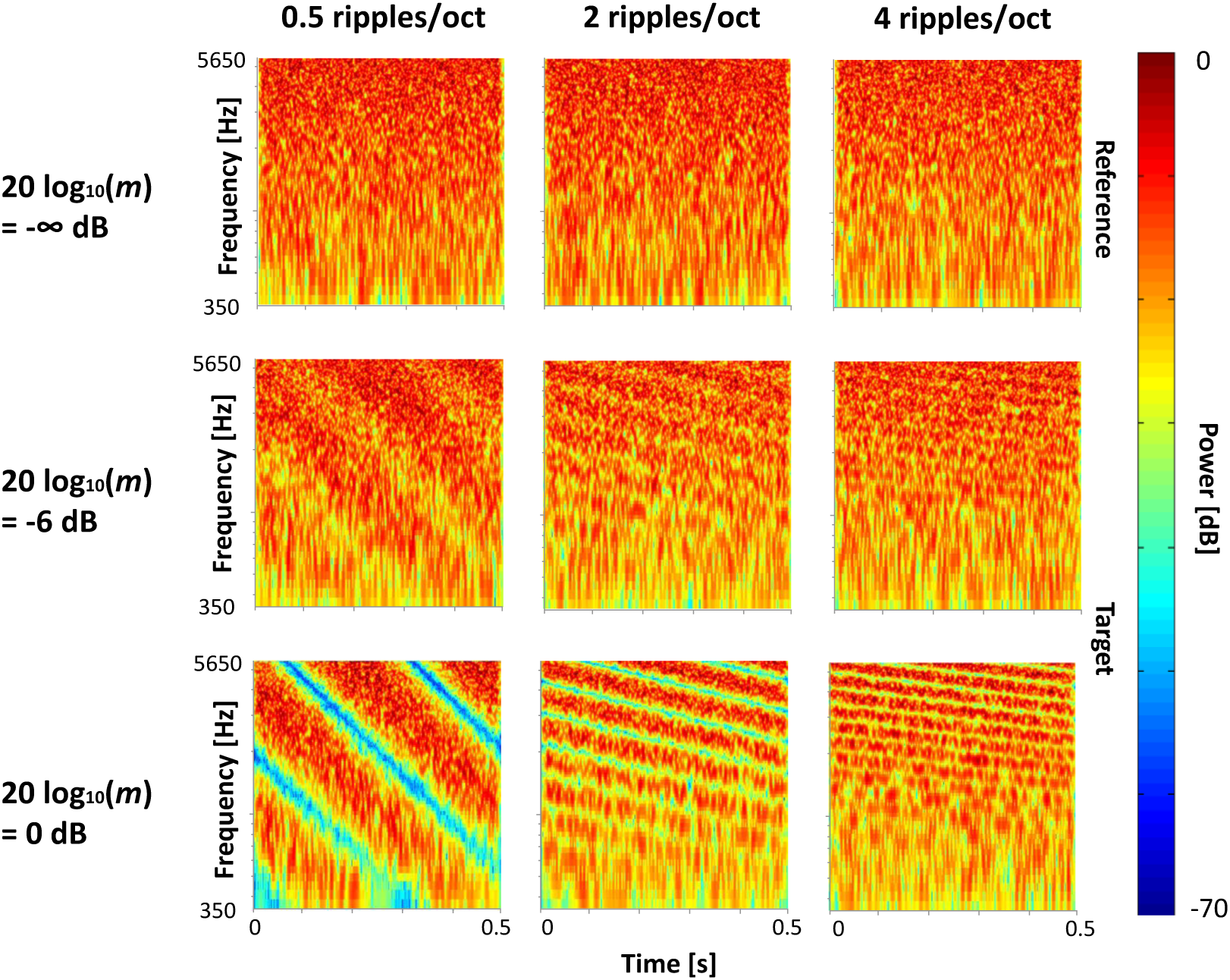
(Color online) Spectrograms of the spectrotemporal modulated stimuli. Notice how the pattern along the frequency axis changes with increasing ripple density.

#### 2.1.4. Procedure

Initially, the stimuli were presented at levels of 65 and 86 dB SPL using all three ripple densities (0.5, 2, and 4 ripples/oct). Then, stimuli were presented at levels of 55, 65, 75, and 86 dB SPL with a ripple density of 4 ripples/oct. Throughout, stimuli were presented monaurally to the left ear. Level roving of 8 dB was used (i.e., random gain between −4 and 4 dB for each stimulus) to reduce the salience of level cues (Eddins and Bero, 2007).

A two-interval two-alternative forced-choice task was used. One of the intervals contained the unmodulated (i.e., reference) stimulus and the other interval contained the modulated (i.e., target) stimulus. The target was randomly presented in the first or second interval with equal probability. There was a 500-ms pause between intervals. Participants were seated in front of a computer screen. They were instructed to discriminate the target interval, which would correspond to the stimulus with a “rippled, vibrating sound”, from the reference interval, which would correspond to the stimulus with a “noisy sound”. They did so by clicking on the corresponding button on the screen (or by using the corresponding keys on the keyboard). Visual feedback was provided through a green (correct response) or red (incorrect response) highlight after each trial. Conditions were presented to each participant in a random order. In a given run, the ripple density was fixed. The modulation depth at threshold was estimated using a three-down one-up procedure tracking the 79.4% point on the psychometric function (Levitt, 1971). Each run started with a fully modulated target (20 log_10_(*m*) = 0 dB). The modulation depth was decreased by 6 dB after the first reversal, changed by 4 dB until two more reversals occurred, and changed by 2 dB for the last 6 reversals. A run was ended after 9 reversals. For each run, the mean value of 20 log_10_ *m* at the last 6 reversals was calculated. Participants completed a test and retest run for every condition. If the thresholds for the two differed by more than 3 dB, a third run was completed. For each condition, the final threshold was taken as the average of all runs.

### 2.2. Results

Statistical analysis was conducted using the R programming language and statistical environment (R Core Team, 2017).

Figure 2 shows a boxplot of the STMD thresholds together with the average across participants for stimuli at 65 and 86 dB SPL and 0.5, 2 and 4 ripples/oct. A general linear model (GLM) showed that ripple density had a significant effect on the STMD thresholds (χ^2^(1) = 8.26, *p* < 0.001) as did level (χ^2^(1) = 11.76, *p* < 0.001). There was a significant interaction of ripple density and level (χ^2^(1) = 24.17, *p* < 0.001). Tukey *post hoc* tests on the GLM revealed increased thresholds with increasing ripple density at 86 dB SPL, between 0.5 ripples/oct and 4 ripples/oct (*z* = 5.83, *p* < 0.001, confidence interval (CI) [3.64, 8.02]) and between 2 ripples/oct and 4 ripples/oct (*z* = 4.82, *p* < 0.001, CI [2.63, 7.01]). In contrast, thresholds decreased with increasing ripple density at 65 dB SPL between 0.5 ripples/oct and 2 ripples/oct (*z* = −3.57, *p* < 0.001, CI [−5.76, −1.38]) and then increased between 2 ripples/oct and 4 ripples/oct (*z* = 2.54, *p* = 0.012, CI [0.35, 4.73]). The STMD thresholds were significantly lower at 65 dB SPL than at 86 dB SPL at 2 ripples/oct (*z* = 3.12, *p* < 0.001, CI [0.93, 5.31]) and at 4 ripples/oct (*z* = 5.40, *p* < 0.001, CI [3.21, 7.59]), but not at 0.5 ripples/oct (*z* = −1.45, *p* = 0.40, CI [−3.64, 0.73]).

**Figure 2:**
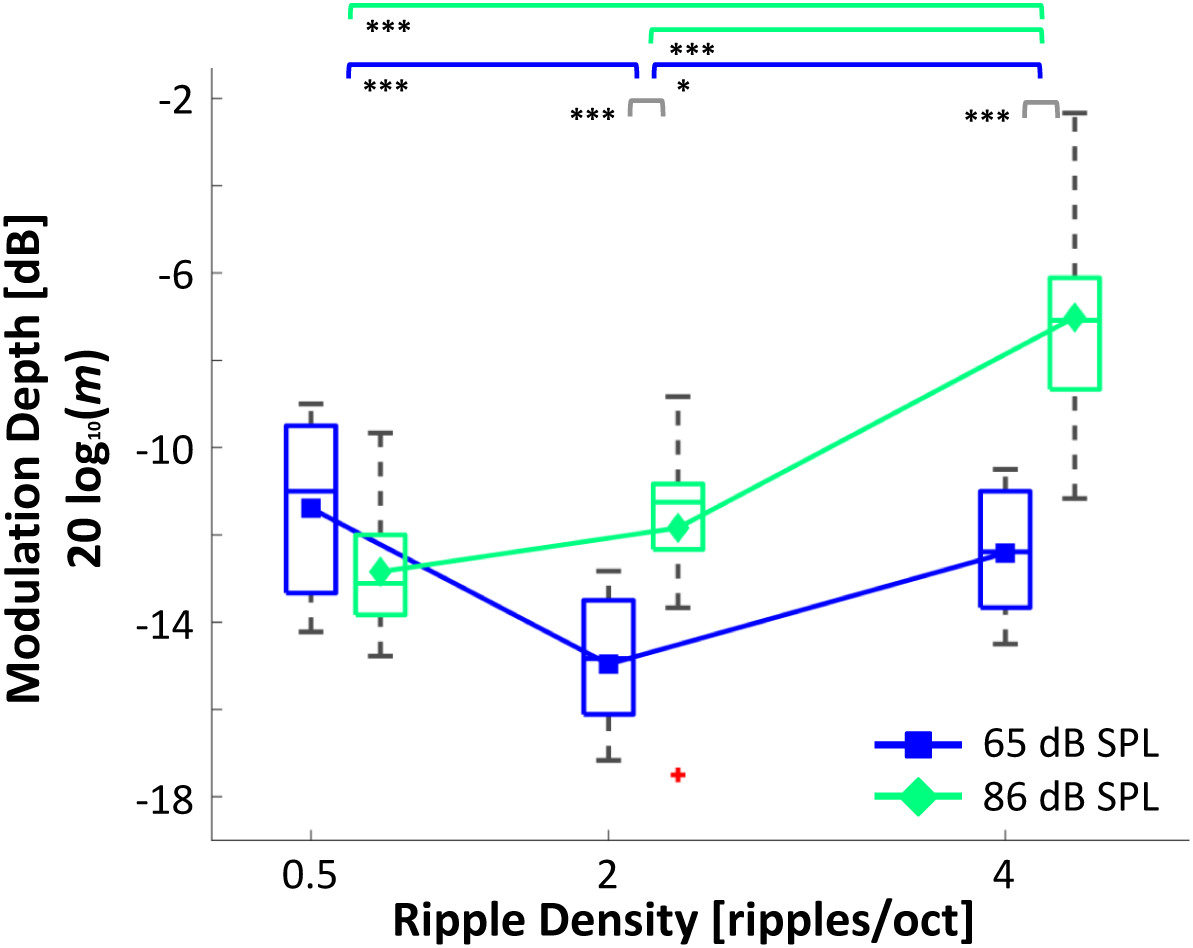
(Color online) STMD thresholds boxplot. The line in the middle of each box represents the median of the participants’ thresholds for each condition. Symbols represent the average. The vertical edges of each box represent the 25th and 75th percentiles. The distance between them is the interquartile range (IQR). Error bars (i.e., whiskers) are drawn from the ends of the IQR to the furthest data point within 1.5 of the IQR. Crosses represent data points beyond that (i.e., outliers). Lower thresholds indicate better performance. * = *p* < 0.05, *** = *p* < 0.001.

Figure 3 shows a boxplot of the STMD thresholds for stimuli at 55, 65, 75, and 86 dB SPL and 4 ripples/oct. STMD thresholds changed significantly with level (Friedman’s ANOVA, 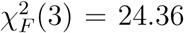, *p* < 0.001). There was a large increase between 65 and 75 dB SPL. *Post hoc* Conover’s tests with Holm correction for multiple comparisons revealed significant differences between 55 and 75 dB SPL (*p* < 0.001), 55 and 86 dB SPL (*p* < 0.001), 65 and 75 dB SPL (*p* < 0.001), 65 and 86 dB SPL (*p* < 0.001), and 75 dB SPL and 86 dB SPL (*p* < 0.001). There was no significant difference between thresholds for the two lowest levels (55 and 65 dB SPL, *p* > 0.05).

**Figure 3:**
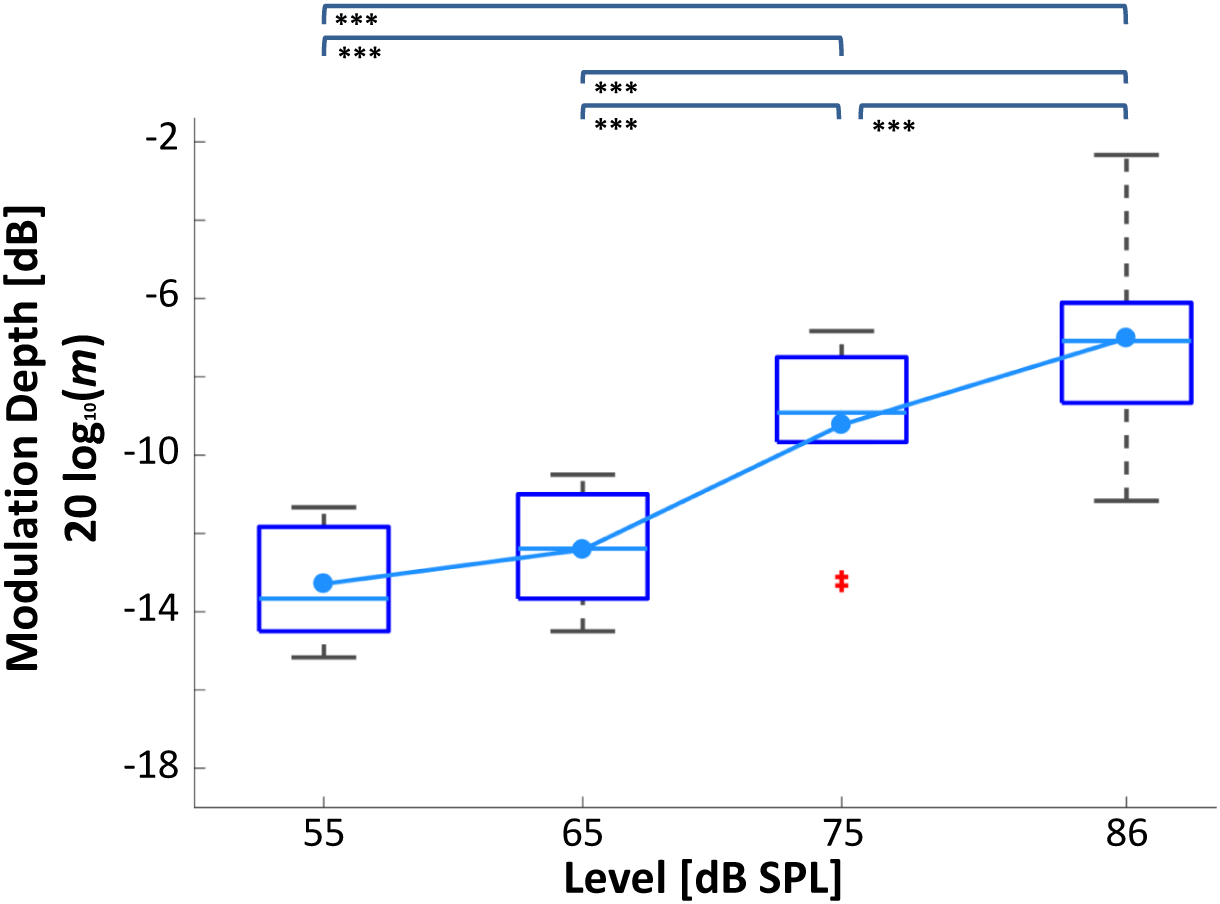
(Color online) STMD thresholds boxplot with a ripple density of 4 ripples/oct. Otherwise as Figure 2.

## 3. Computational Model

We used a computational model with a physiologically inspired front end (i.e., model of the auditory periphery up to the AN) to help us interpret the behavioral results, to study the contribution of peripheral information to spectrotemporal sensitivity, and to obtain quantitative predictions of the behavioral thresholds. A block diagram of the model is shown in Fig. 4. We hypothesized that the model would reflect a worsening in the spectrotemporal representation in the AN with increasing level.

**Figure 4:**
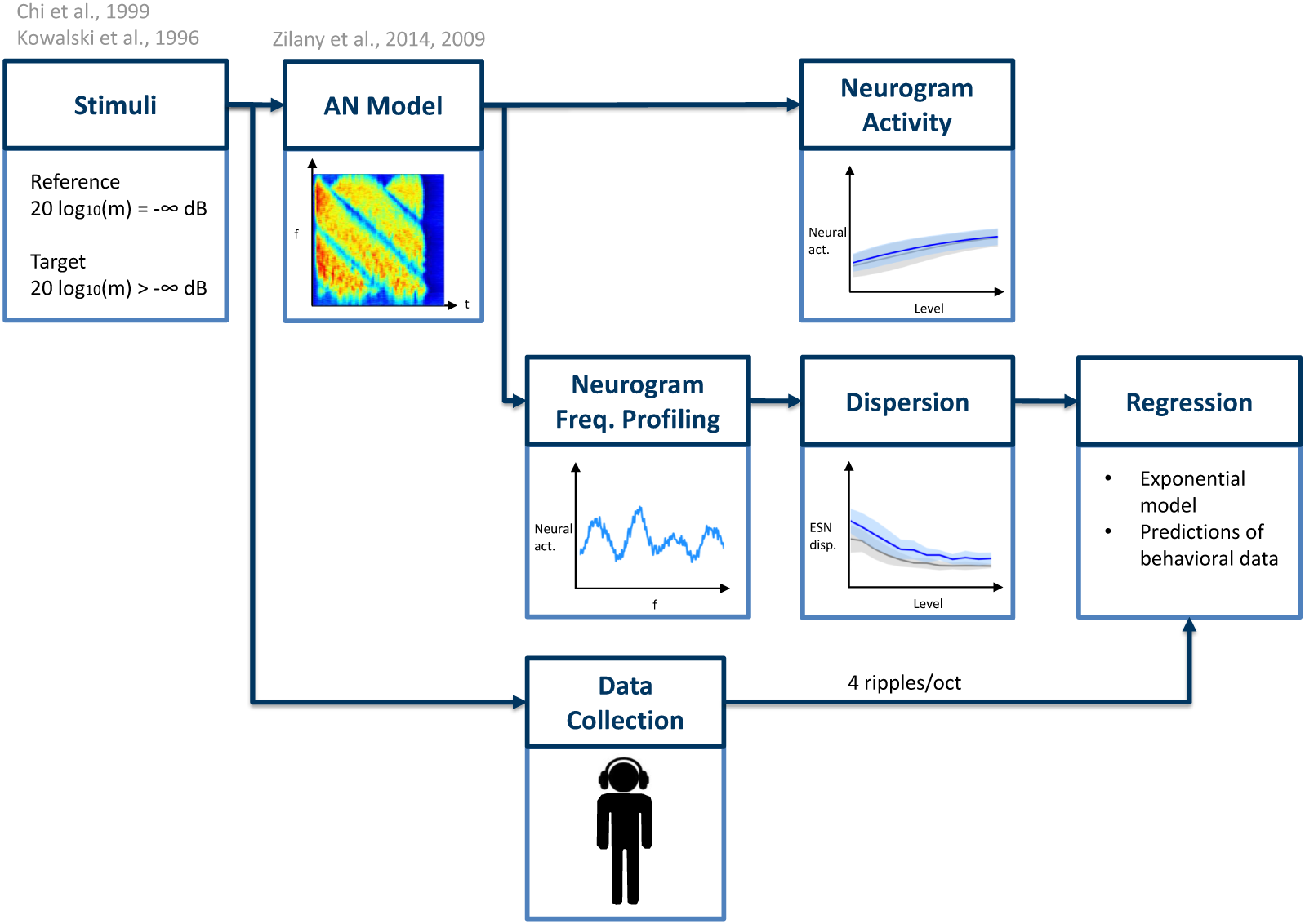
(Color online) Block diagram of the computational model used to interpret the behavioral data and to study the contribution of peripheral information to spectrotemporal sensitivity.

### 3.1. Stimuli

We included a wider range of levels (from 40 to 95 dB SPL in steps of 5 dB) than used in the experiments. We used the same ripple densities (0.5, 2, and 4 ripples/oct). We simulated responses to the reference stimulus (20 log_10_(*m*) = −∞ dB) and target stimuli with a modulation depth of 20 log_10_(*m*) = −6 dB and 20 log_10_(*m*) = 0 dB (which correspond to 50 and 100% modulation, respectively, in a linear scale).

### 3.2. AN Model

The model proposed by Zilany et al. (2009, 2014) was used as a front end. This model reproduces the responses of AN fibers to acoustic stimulation. It has been validated with a wide range of physiological data. It is comprised of different modules (each simulating a specific function of the auditory periphery).

First, the stimulus is passed through a filter simulating the middle ear frequency response. The output is fed to a signal path and a control path. The signal path mimics the behavior of the outer-hair-cell-(OHC-) controlled filtering of the basilar membrane in the cochlea and the transduction of the inner-hair-cells (IHCs) by a series of non-linear and low-pass filters. The control path mimics the function of the OHCs in controlling basilar membrane filtering. The control path output feeds back into itself and into the signal path. The output of the IHCs is fed to the IHC-AN synapse module with two power-law adaptation paths, which simulate slow and fast adaptation.

For each stimulus, the AN model generated a so-called early stage neurogram (ESN). An ESN is a time-frequency representation of a signal which encodes temporal modulations caused by the interaction of spectral components in each band (Elhilali et al., 2003). It shows the response of neurons tuned to different characteristic frequencies (CFs) through time. We used 512 CFs logarithmically spaced from 250 to 8000 Hz. For each CF, we simulated the average response of 50 AN fibers with different spontaneous rates: high (100 spikes/s), medium (5 spikes/s), and low (0.1 spikes/s), with proportions of 0.6, 0.2, and 0.2, respectively, which correspond to the distribution observed in mammals (Liberman, 1978; Zilany and Bruce, 2007). We grouped the neural activity into time bins of 8 ms, which is close to the equivalent rectangular duration of the temporal window of the auditory system (Moore et al., 1988; Oxenham and Moore, 1994). Afterwards, we smoothed the response by convolving it with a 2-sample long rectangular window with 50% overlap. Figure 5 shows example ESNs of reference and target stimuli for different ripple densities.

**Figure 5:**
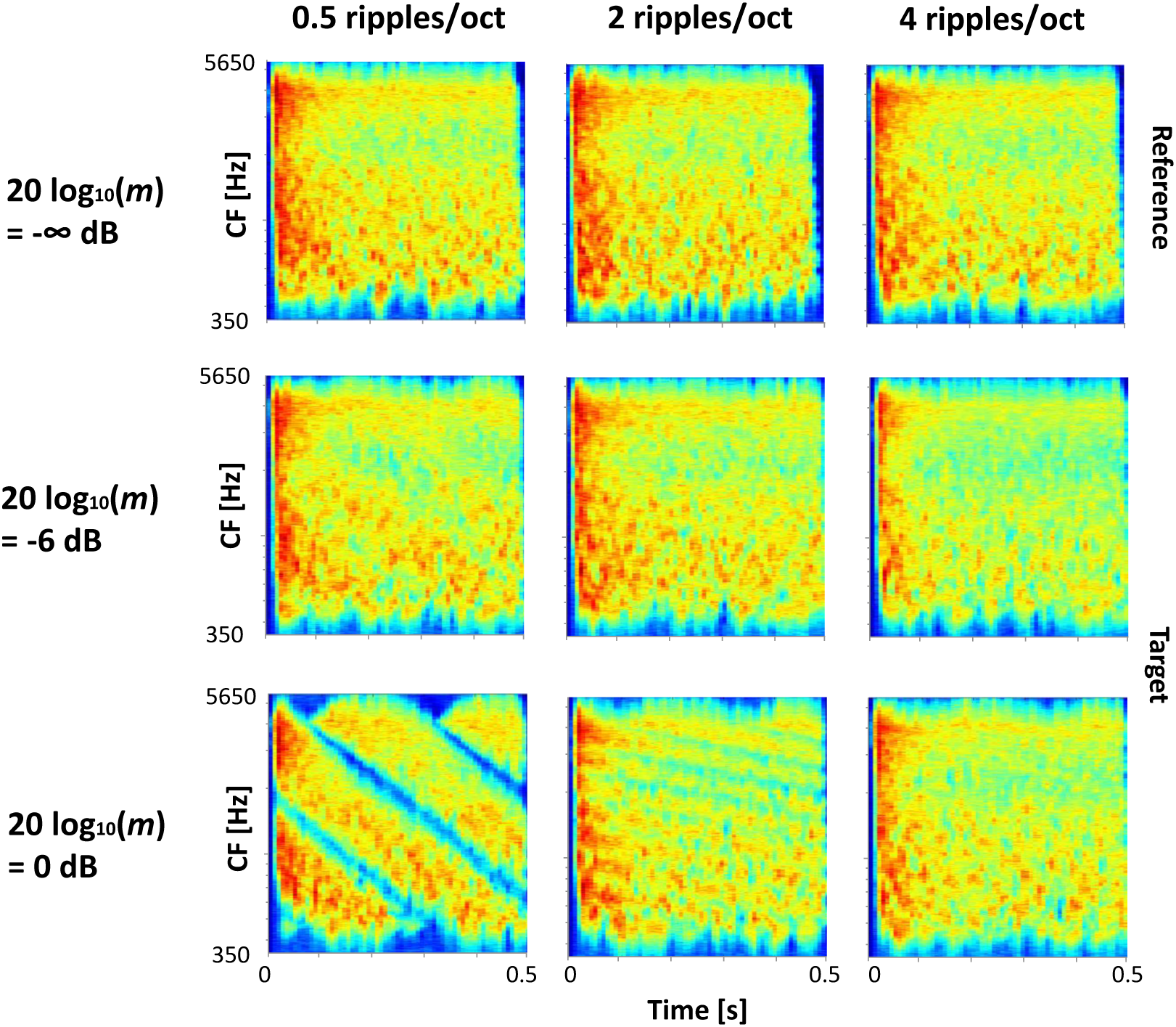
(Color online) ESNs of spectrotemporally modulated stimuli.

### 3.3. Neurogram activity

We quantified the increase of neural activity by computing the mean and standard deviation of the neurograms across different levels. Figure 6 shows plots of the ESN activity for the reference stimulus (20 log_10_(*m*) = −∞ dB) and a fully-modulated target stimulus (20 log_10_(*m*) = 0 dB). In all cases, increasing the level increased the neural activity and the slope of the curves decreased at high levels. These trends were consistent across all three ripple densities.

**Figure 6:**
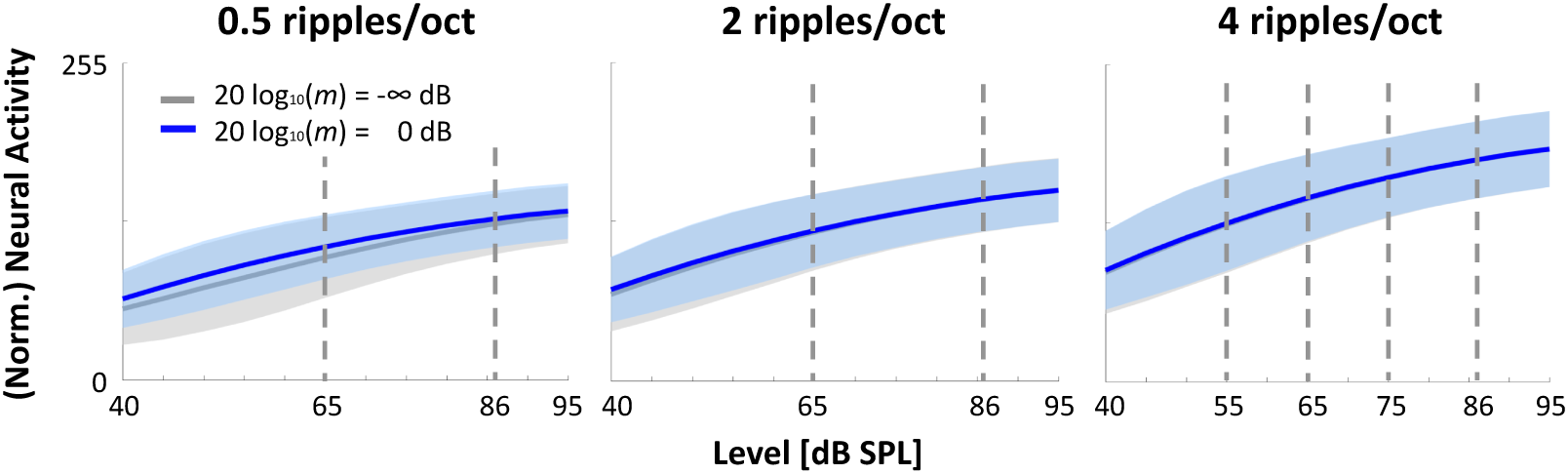
(Color online) Neurogram activity. The solid lines and the shaded areas correspond to the mean and standard deviation, respectively, of each neurogram. This is a measure of the amount of activity at the AN level. A large increase in activity could lead to saturation and, therefore, to a poorer spectrotemporal representation, yielding higher thresholds (Sec. 4). The dashed lines represent the levels at which behavioral measurements were obtained.

### 3.4. Neurogram frequency profiling

We defined a *frequency profile* of a neurogram as a slice across its CFs at a given point in time. If we think of a neurogram as an image, a frequency profile would correspond to all the row values of a specific column.

Consider the ESNs in Fig. 5 for the ripple density of 0.5 ripples/oct. The top ESN (20 log_10_(*m*) = −∞ dB) shows a uniform, indistinct pattern. A frequency profile at any point in time would show a roughly flat curve. In contrast, the bottom ESN (20 log_10_*(*m*)* = 0 dB) shows a clear pattern, reflecting the spectrotemporal characteristics of the stimulus. A frequency profile at any point in time would show distinct crests and troughs. Figure 7 shows frequency profiles for different ripple densities at different levels.

**Figure 7:**
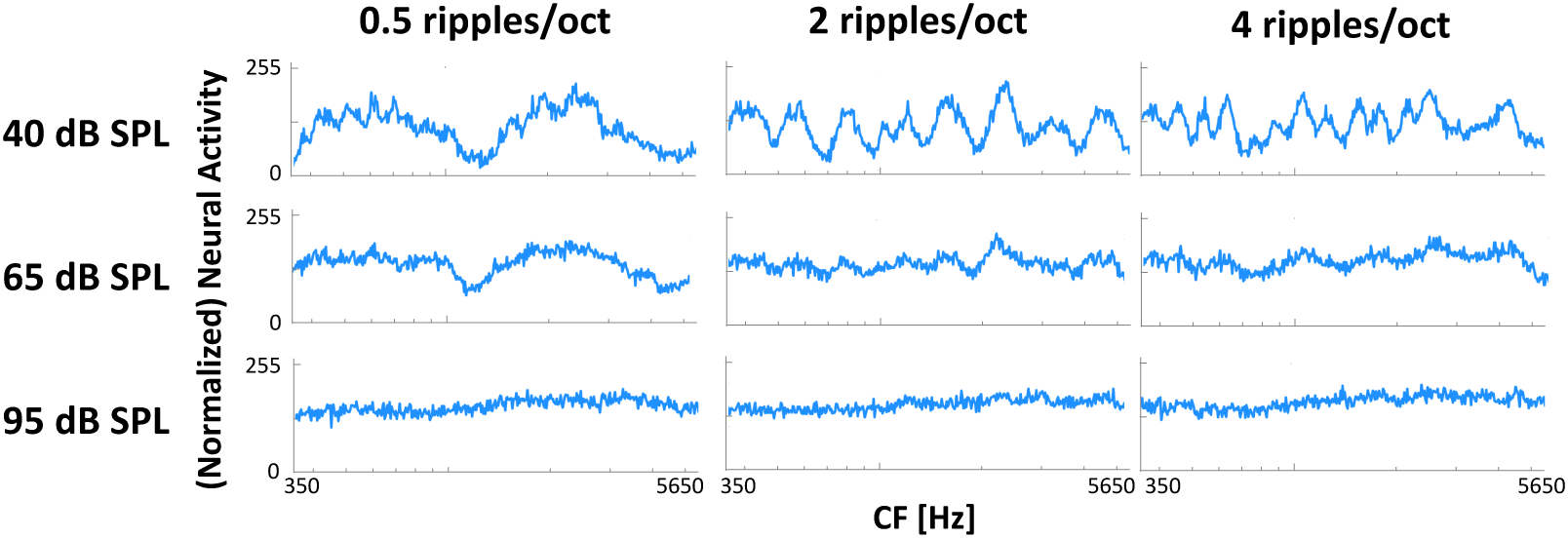
(Color online) Frequency profiles of the ESNs at *t* = 250 ms (total duration of the stimulus was 500 ms).

### 3.5. Dispersion

One measure of the information available at the AN level for detection of modulation is the dispersion of the frequency profiles (i.e., columns) of the ESNs across time. The dispersion is a measure of the amplitude of the frequency profile curves. It measures the amount of variation in amplitude across the frequency range. We quantified this dispersion using the interquartile range (IQR), as shown in Eq. 3. We also computed a measure of the dispersion variability across all the frequency profiles of a given neurogram, as shown in Eq. 4. In both cases, ESNj is the frequency profile at the *j*-th point in time.

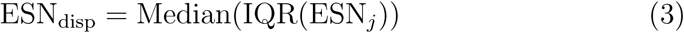

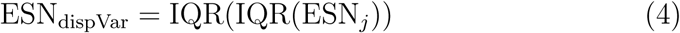

Figure 8 shows plots of ESN dispersion. Deeper modulations (closer to 20 log_10_ (*m*) = 0 dB) led to larger dispersions for lower ripple densities (0.5 and 2 ripples/oct). In all cases, increasing the level reduced the dispersion. This trend was consistent across all three ripple densities.

**Figure 8:**
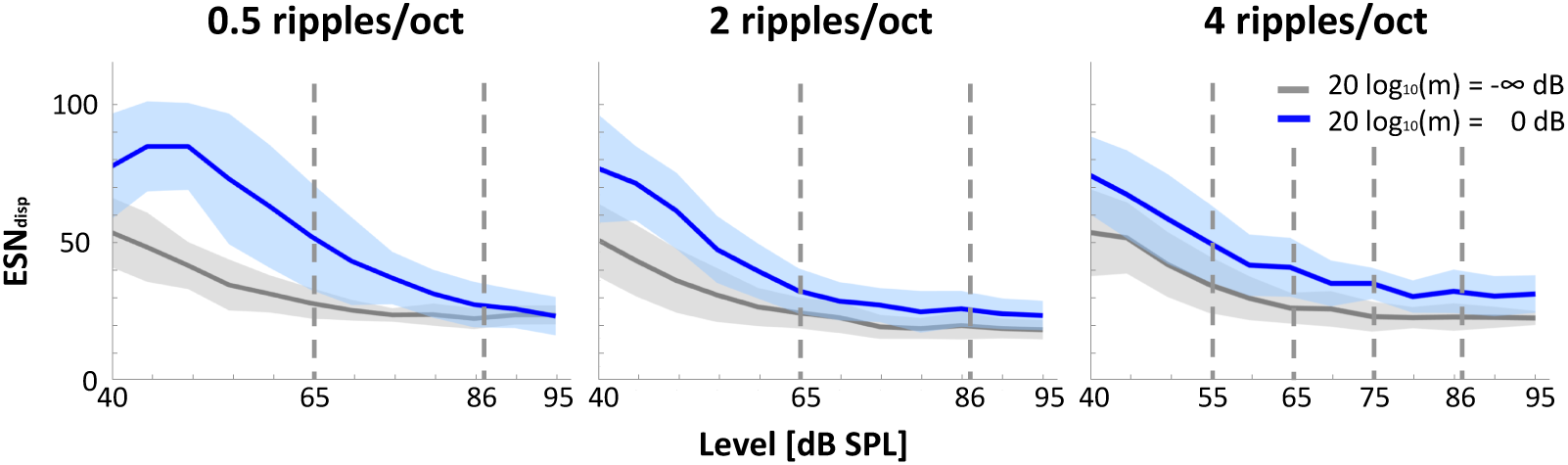
(Color online) Plots of ESN dispersion. The solid lines and the shaded areas correspond to the ESN dispersion and dispersion variability, respectively, for frequency profiles across all time points for each neurogram. The ESN dispersion is a measure of the amount of information for modulation detection available at the AN level (larger dispersion allows for higher detectability, Sec. 4). The dashed lines represent the levels at which behavioral measurements were obtained.

### 3.6. Regression

Model results were compared with the behavioral data using a regression model. Since the results of experiment 1 showed that the effect of presentation level was largest at 4 ripples/oct, we focused on the behavioral data for experiment 2.

We calculated the difference in dispersion between a fully-modulated target stimulus (20 log_10_(*m*) = 0 dB) and the non-modulated reference as a predictor for an exponential regression model as described by Eq. 5:

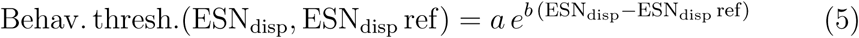

with parameters *a* and *b*. It yielded an (adjusted) *R*^2^ value of 0.98 and a root mean squared error (RMSE) of 0.25 dB. Figure 9 shows plots of the behavioral data versus the model metric as well as the regression model. We used the generated model to predict the behavioral thresholds for the different levels. Figure 10 shows the model’s predictions as well as the mean of the behavioral data (as a reference). The model predictions show that the lowest (best) STMD threshold is around 20 log_10_(*m*) = −13.5 dB for the modelled experiment.

**Figure 9:**
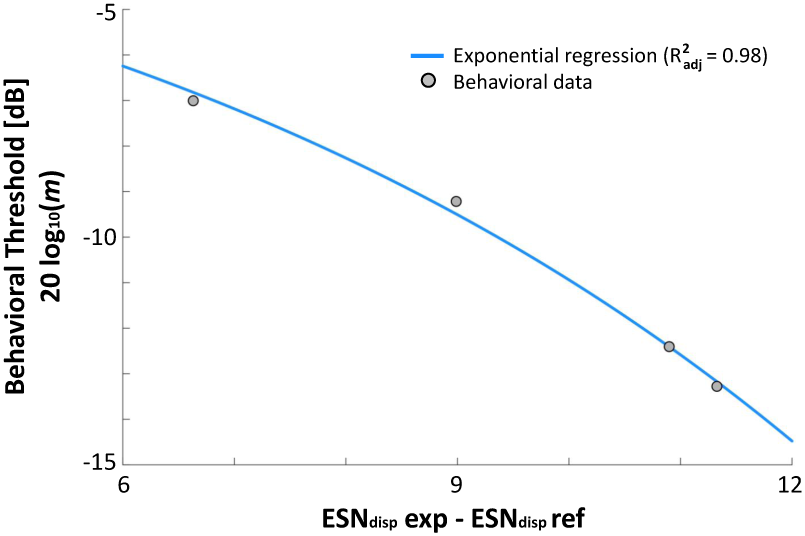
(Color online) Exponential regression model of the behavioral thresholds of experiment 2 (ripple density of 4 ripples/oct).

**Figure 10:**
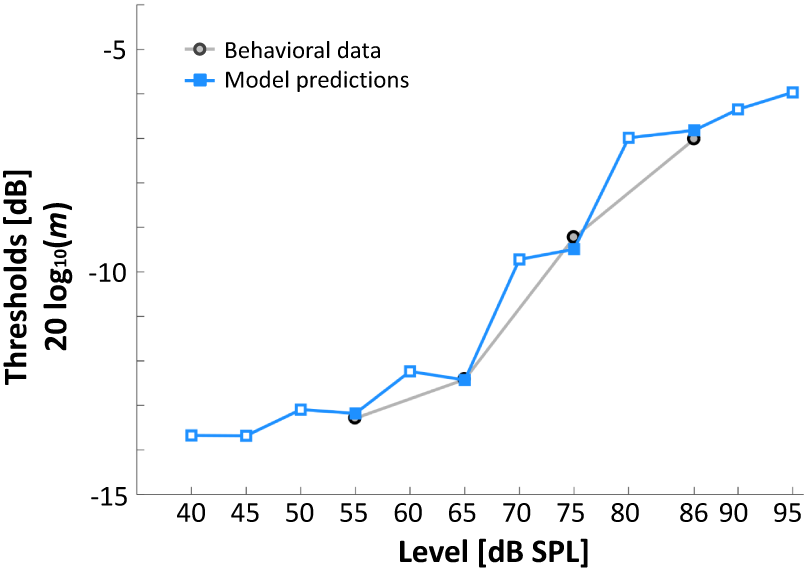
(Color online) Model predictions (squares) of the behavioral thresholds (circles) across different levels (ripple density of 4 ripples/oct).

## 4. Discussion

Higher levels led to increased STMD thresholds. Moreover, increasing ripple density affected the STMD thresholds differently depending on the level. At 65 dB SPL, STMD thresholds were lowest at 2 ripples/oct. In other studies a similar trend was found. Anderson et al. (2012) found lowest thresholds at 3 ripples/oct, followed by increasing thresholds with increasing ripple density (up to 64 ripples/oct). The participants of Eddins and Bero (2007) performed best either at 2 or 3 ripples/oct. Davies-Venn et al. (2015) found a significant improvement in thresholds from 0.5 to 1 and from 1 to 2 ripples/oct. Other studies (Bernstein and Green, 1987; Leek and Summers, 1996) have found similar trends. The most common explanation is that there are two regions in which different cues are used. For low ripple densities (<= 3 ripples/oct), the ripples are detected using a spectral-contrast mechanism, while for higher ripple densities (> 3 ripples/oct), the spectral cues become weaker and the interaction between the close peaks in the rippled noise provides usable temporal cues (Davies-Venn et al., 2015). However, further studies are needed to confirm this. At 86 dB SPL, STMD thresholds increased with increasing ripple density, similar to what Bernstein et al. (2013) found. The effect of presentation level was largest at 4 ripples/oct, where low presentation levels (55 and 65 dB SPL) yielded significantly lower (better) STMD thresholds than high presentation levels (75 and 86 dB SPL).

Understanding the effect of level on STMD thresholds for NH listeners is the first step to understanding it in HI listeners. Although it is very likely that level also affects STMD thresholds of HI listeners, our results cannot be translated directly to the HI population for several reasons. Firstly, increasing the intensity affects neural saturation for NH and HI listeners differently. This can also affect perception differently due to the abnormal loudness-growth curve (i.e., non-linear loudness shift, Edwards et al., 1998; Hellman, 1999) of HI listeners. Additionally, the auditory filters of HI listeners are abnormally broad (resulting in spectral smearing of the internal representation of the stimulus, Moore, 2007) and change less with level compared to NH listeners. Furthermore, the large heterogeneity of the HI population (Lopez-Poveda and Johannesen, 2012) would very likely play a role. Therefore, we hypothesize that STMD thresholds of HI listeners will also be affected by level and will be worse than those of NH listeners. However, testing this requires further behavioral measurements and modelling. This would be a crucial step for further understanding the differences in STMD thresholds between NH and HI participants. Our results show that attributing them to differences in spectrotemporal sensitivity would be only partially true, since level also plays an important role.

We used a computational model with a physiologically inspired front end to explain the behavioral results (Fig. 5). We found that the observed effects of level on the behavioral data could be explained by a worsening of the spectrotemporal representation in the AN due to broadening of the cochlear filters. Furthermore, higher levels led to more neural saturation “filling in the dips” of the neurograms. This can be seen in the increase of the neural activity (Fig. 6) and the flattening of the frequency profiles (Fig. 7). Frequency profiles at lower levels reflected the changes of the spectral information across time, while frequency profiles at higher levels lost the representation of this information (Fig. 8). All these factors diminish the coding of the spectrotemporal pattern of the modulated stimuli in the AN with increasing level, making it harder to discriminate.

The regression analysis (Fig. 9) suggested that information in the auditory periphery is able to account for a large proportion of the variance in the behavioral data, supporting its value for predicting spectrotemporal modulation thresholds (Fig. 10).

Similar results could have been obtained with a more simple model (e.g., an excitation pattern model, Moore and Glasberg, 1987). However, the use of frameworks based on the biology of the auditory system has a few advantages. For instance, they incorporate physiological information inherently. This allows a more direct, transparent understanding of the auditory mechanisms at different stages of the auditory pathway (the periphery in this case), since it gives insight into the representation of the stimuli at each of these steps. Additionally, the Zilany et al. (2009, 2014) AN model incorporates the effects of sensorineural hearing loss due to damage to the IHCs and OHCs (something that would not be straightforward to do using a nonphysiological approach). Now that presented framework has been validated for the NH case, this would be of special interest, since it could allow studying the effect of level on spectrotemporal modulation detection by HI listeners using a similar framework to the one described here.

Furthermore, alternative back ends could have been used in the proposed model. For example, the ratio between the dispersion of the reference and the target stimulus (instead of the difference) could have been used as the predictor for the regression. Additionally, a different approach could have been used to predict the behavioral threshold. For instance, the difference in dispersion (or the quotient) between the reference and the target stimulus required for threshold could be computed. Afterwards, the modulation depth required to achieve this difference metric could be calculated iteratively, with the final value being the predicted behavioral threshold. This approach would eliminate the need for the regression model in Eq. 5.

The effect of presentation level has a number of implications for the use of STMD tests in experimental and clinical environments. When administering STMD tests at different levels, the observed differences in STMD thresholds should (at least partially) be attributed to the effect of level, making it more complex to interpret the contribution of spectrotemporal sensitivity only. For NH participants it is recommended to use a fixed presentation level to allow for direct comparison between their STMD thresholds. However, it is unclear how level affects STMD thresholds in HI listeners. Therefore recommendations for STMD tests in HI participants cannot be made based on our data. Future work will be focused on investigating level effects for different types of spectral and spectrotemporal ripple tests, as well as for HI listeners.

## 5. Conclusions

STMD thresholds were higher (worse) at high than at low presentation levels, with larger differences in thresholds at 4 ripples/oct than at 2 ripples/oct. The computational model with a physiologically inspired front end could account for the behavioral results, showing that information at the peripheral level is sufficient to predict the behavioral thresholds. STMD thresholds obtained at different presentation levels are affected not only by differences in spectrotemporal modulation, but also at least partly by level. Therefore, this effect needs to be considered when administering STMD tests (both in clinical practice and in experimental research) and when comparing STMD thresholds within and across studies.

## Acknowledgements

This work was supported by a TBM-FWO grant from the Research Foundation-Flanders [grant number T002216N] and an ERC Starting Grant (to Tom Francart) from the European Research Council under the European Union’s Horizon 2020 Research and Innovation Programme [grant agreement no. 637424]. In addition, this research was jointly funded by Cochlear Ltd. and Flanders Innovation & Entrepreneurship (formerly IWT), project 50432. We would like to thank Benjamin Dieudonné for his technical help generating the stimuli. We would also like to thank the editor Dr. Brian C. J. Moore and the three anonymous reviewers for their helpful comments and feedback. Finally, our most sincere thanks to our participants for their time and dedicated effort.

